# The kinesin-8 member Kif19 alters microtubule dynamics, suppresses cell adhesion, and promotes cancer cell invasion

**DOI:** 10.1101/2020.09.04.282657

**Authors:** Samuel C. Eisenberg, Abhinav Dey, Rayna Birnbaum, David J. Sharp

## Abstract

Metastasis is one of the deadliest aspects of cancer. Initial Metastatic spread is dependent on the detachment and dissemination of cells from a parent tumor, and invasion into the surrounding tissue. In this study, we characterize the kinesin-8 member Kif19 as a promoter of cancer cell invasion that suppresses cell-cell adherens junctions and cell-matrix focal adhesions. Initial analysis of publicly available cancer patient data sets demonstrated that Kif19 expression correlates with worse overall survival probability in several cancers and that Kif19 expression is increased in metastases of colorectal and breast carcinoma compared to the primary tumor. Depletion of Kif19 from two human cancer cell lines (DMS53 and MDA-MB-231) did not alter viability, but decreased the cells’ ability to invade a Matrigel matrix by half and impaired the invasion of spheroids into a primary cell monolayer. Ectopically expressed Kif19 localized to, and partially depolymerized, microtubules in the cell periphery. However, Kif19 depletion increased microtubule dynamicity and sensitivity to pharmacological depolymerization without altering total microtubule polymer levels. These data indicate that Kif19 can both depolymerize and stabilize microtubules. Given this activity, we then studied Kif19’s effect on focal adhesions and adherens junctions, which are both regulated by microtubule dynamics. Kif19 knockdown increased the proportion of cell surface area covered by Vinculin focal adhesions. Further, Kif19 depletion increased whole cell E-cadherin expression and the accumulation of E-cadherin at cell-cell adherens junctions. Conversely, ectopic overexpression of full-length Kif19 led to proportionally smaller focal adhesions and impaired E-cadherin accumulation at cell-cell junctions. Our current hypothesis is that aberrant Kif19 expression in cancer alters focal adhesion dynamics and suppresses E-cadherin expression, which enhance cell invasiveness. Further, we propose that these changes in cell adhesion are due to modification of peripheral microtubule dynamics by Kif19, potentially through disruption of local rho GTPase activity.

## Introduction

Metastasis is one of the primary mechanisms by which cancer causes mortality. More than 90% of cancer-related deaths are attributable to metastatic disease(1, 2). Metastasis can be thought of as a stepwise derangement in cellular function, facilitating first escape from the parent tumor site, then migration to a new area, invasion of the host tissue, and finally successful growth in the new environment (1). Significant work has been to done to understand the mechanisms underlying dissociation from and migration away from the parent tumor, as anti-metastatic therapeutics properties could greatly improve cancer prognosis (2, 3). An early step in cancer metastasis, known as the epithelial mesenchymal transition (EMT), has been studied extensively as a marker for metastatic potential (4, 5). The EMT constitutes a global change in protein expression during which cells lose epithelial characteristics in favor of mesenchymal characteristics. This results in cells that are more motile, invasive, and resilient (5). Recent work has indicated that EMT is not a binary event, as metastatic cells can exhibit a range of epithelial and mesenchymal characteristics depending on the environment (6, 7). However, there are some characteristics of early EMT and invasion that are essential for the initiation of metastasis in multiple cancer types, specifically dissociation and movement away from the parent tumor (8, 9). Two major mediators of these processes are focal adhesions (FAs) and adherens junctions (AJs), which mediate cell-extracellular matrix (ECM) and cell-cell adhesions, respectively (7, 10, 11). The partial or total loss of E-cadherin, a major component of AJs in epithelial-like cells, is often considered a marker of metastatic potential (12-14). Multiple FA components, including integrins, focal adhesion kinase (FAK), and Paxillin are associated with cancer aggressiveness and invasiveness (15-17). Thus, the regulation of FA and AJ formation, maturation, and disassembly are important topics for better understanding cancer metastasis and invasion (7, 10, 16).

AJs and FAs share several common regulatory mechanisms. Both structures are bound to the actin cytoskeleton and share many of the components mediating this attachment, namely Vinculin, α-Actinin, Zyxin, and VASP (18-21). The Rho GTPases regulate AJs and FAs via actin dynamics. Rac1-mediated branched-actin polymerization has been implicated in the formation of both FAs and AJs, while RhoA-mediated actin stress fiber formation and myosin II contractility are implicated in the maturation and stabilization of both structures (22-25). Lastly, both systems can be regulated by the microtubule cytoskeleton.

Microtubules (MTs) are dynamic cytoskeletal polymers composed of α- and β-tubulin heterodimers (26). They exhibit “dynamic instability,” a propensity to vacillate between periods of growth and shrinkage. The rapid transition from growth to shrinkage is termed catastrophe and the inverse is termed rescue (27). MTs are classically associated with cell division, ciliogenesis, and axon maintenance (28-30). However, there is a growing body of data implicating MTs in the regulation of cell movement and adhesion (31-35). MT dynamics in the cell are highly regulated by stabilizing and destabilizing factors (31, 36-39). Kinesins are a protein family that have been identified as important regulators of MT dynamics in recent years (40-42). These are classically dimeric ATPase motor proteins that move along MTs and transport proteins, vesicles, or RNA to various parts of the cell (43, 44). Isolated members from several kinesin families have been associated with MT stabilization, but this is often a secondary function of the protein (45-47). However, two kinesin families, kinesin-8 and kinesin-13, are specifically associated with MT dynamics, typically destabilization (40). Members of the kinesin-8 family are notable for containing a MT plus-end directed motor that can additionally depolymerize MT plus-ends (48). However, full-length kinesin-8 proteins are reported to have additional effects at MT plus-ends, including both decreasing and increasing MT dynamics (49-51). This variation in function is partly determined by the kinesin-8 C-terminal tail (CTT), which regulates the protein localization and activity (52-54). There are three known kinesin-8 family members in mammals (48, 55). Two of these, Kif18A and Kif18B, have recently been identified as potential oncogenes, and their upregulation has been associated with worse outcomes in several cancers (56-63). The third mammalian kinesin-8 member, Kif19, is relatively uncharacterized, though it has recently been implicated as a peripheral blood biomarker for cervical cancer (64-66). The Kif19 motor domain is capable of both MT plus-end motility and MT depolymerization on purified MTs, similar to other kinesin-8 members (48, 65). Yet while the majority of kinesin-8s localize to the nucleus and regulate mitosis, Kif19 has been reported to localize to the tips of cilia in mice, where it regulates ciliary length (64, 67-71).

Here, we present data implicating Kif19 as a potential oncogene that regulates cell-cell and cell-ECM adhesion by disrupting MT dynamics at the cell edge. A preliminary search of mRNA expression data in publicly available cancer genomics data sets implies that higher Kif19 expression is associated with significantly reduced overall survival probability in multiple cancer types and that Kif19 expression is greater in metastases from colorectal and breast carcinoma than in the primary tumor. Further, siRNA depletion of Kif19 from small cell lung cancer (DMS53) and breast cancer (MDA-MB-231) cell lines significantly impairs the cells’ ability to invade through a Matrigel matrix. DMS53 spheroids seeded onto a monolayer of primary endothelial cells also show significantly reduced sprouting and invasion following Kif19 knockdown. We show that plasmid-expressed Kif19 localizes to MTs at the cell periphery and is capable of depolymerizing intracellular MTs. However, we also report that Kif19 exhibits MT stabilizing activity: following Kif19 knockdown, there is an increase in the proportion of MTs that are tyrosinated and no associated change in total polymerized MTs, implying a more dynamic MT population. Additionally, we report that the MT array is more sensitive to low-dose nocodazole after Kif19 siRNA knockdown. Thus, Kif19 appears to exhibit both MT-stabilizing and -destabilizing behavior. Immunofluorescent (IF) analysis indicates that cells depleted of Kif19 by siRNA exhibit an increase in FA density, particularly at the cell periphery, while FA density is lower in cells overexpressing plasmid-based Kif19 constructs. Lastly, we show that cells treated with Kif19 siRNA have increased E-cadherin levels at cell-cell contacts and in the whole cell, while cells overexpressing plasmid-based Kif19 have lower E-cadherin levels at cell-cell contacts.

## Materials and methods

### Kaplan Meier survival curves and gene expression analysis using the R2 software

We explored the correlation between Kif19 expression and survival of cancer patients through the R2 application (http://r2.amc.nl) using the log-rank method in multiple datasets, including Tumor Kidney Renal Clear Cell Carcinoma-TCGA-533 patients-(**R2 internal identifier:** ps_avgpres_tcgakirc533_tcgars); Tumor Kidney Renal Papillary Cell Carcinoma-TCGA-290 patients-(**R2 internal identifier:** ps_avgpres_tcgakirp290_tcgars); Tumor Stomach Adenocarcinoma-TCGA-415 patients-(**R2 internal identifier:** ps_avgpres_tcgastad415_tcgars); Tumor Breast (MDC) - Bertucci - 266 patients-(**R2 internal identifier:** ps_avgpres_gse21653geofrma266_u133p2); Tumor Cervical Squamous Cell Carcinoma - TCGA - 305 patients-(**R2 internal identifier:** ps_avgpres_tcgacesc305_tcgars); Tumor Ewing Sarcoma (Core Exon) - Dirksen - 85 patients-(**R2 internal identifier:** ps_avgpres_gse63157geop85_huex10p). The cut-off was identified by methods described in the R2 web-based application (http://r2.amc.nl) (72).

The RNA expression analysis has been done with Megasampler, which is a R2 module for investigating the expression level of a gene in any number of the numerous datasets stored in the R2 database. In this analysis, we compared primary breast tumor RNA expression data (Tumor Breast (HER2) - Concha - 66 - MAS5.0 - u133p2-(**R2 internal identifier:** ps_avgpres_gse29431geo66_u133p2)) with another dataset of RNA expression data for 58 breast cancer metastases from different organs (Tumor Lung Metastases - Massague - 29 - MAS5.0 - u133p2-(**R2 internal identifier:** ps_avgpres_gse14017geo29_u133p2)). The Megasampler analysis for Kif19 expression was also run on two colorectal cancer datasets (Primary Vs Metastatic), Primary Tumor Colon (KRAS mut) - Hase - 59 - MAS5.0 - u133p2 and Metatstatic Tumor Colon CRC cell pop. - Calon - 24 - MAS5.0 - u133p2.

### Commercial antibodies

The commercial antibodies used in these studies are listed below. Parentheses indicate the dilutions used for immunofluorescence (IF) or western blot (WB). Mouse anti-α-tubulin (Thermo Fisher Scientific, MS581P1, IF – 1:600), rabbit anti-α-tubulin (Abcam, ab18251, IF – 1:800), mouse anti-tyrosinated-tubulin (Sigma Aldrich, T9028, IF – 1:600), rabbit anti-detyrosinated-tubulin (Abcam, ab48389, IF – 1:600), mouse anti-E-cadherin (Abcam, HECD-1, IF – 1:600), mouse anti-Vinculin (Novus Biologicals, NB600-1293, IF – 1:400), chicken anti-Myc (Novus Biologicals, NB600-334, IF – 1:600, WB – 1:2000), rabbit anti-Kif19 AV34084 (Sigma, AV34084, WB – 1:1500), rabbit anti-Kif19 HPA073303 (Sigma Aldrich, HPA073303, WB – 1:1500), rabbit anti-RFP (Rockland, 600-401-379, IF – 1:800, WB – 1:2000), mouse anti-GAPDH (Fitzgerald, 10R-G109A, WB – 1:25000), anti-Mouse-HRP (Abcam, WB – 1:5000), anti-Rabbit-HRP (Abcam, WB – 1:5000), anti-Chicken-HRP (Invitrogen, WB – 1:5000).

The following fluorescent secondary antibodies were used for immunofluorescent imaging. They were acquired from Thermo Fisher Scientific and used at a dilution of 1:800. Anti-mouse-Alexafluor488, anti-mouse-Alexafluor568, anti-mouse-Alexafluor647, anti-rabbit-Alexafluor488, anti-rabbit-Alexafluor555, anti-rabbit-Alexafluor568, anti-rabbit-Alexafluor647, anti-chicken-Alexafluor488, anti-chicken-Alexafluor568, anti-chicken-Alexafluor647. Actin was stained with phalloidin-Alexafluor568 (Thermo Fisher Scientific) or phalloidin-Alexafluor488 (Thermo Fisher Scientific).

### Plasmid construction

PCR was conducted using Q5 High-Fidelity DNA Polymerase (New England Biolabs, M0491) according to manufacturer protocol with 400-800ng template DNA, and all restriction enzymes were purchased from New England Biolabs. Primers were generated using Geneious 9.X software and purchased from Integrated DNA Technologies. PCR and restriction digest products were purified using agarose gel electrophoresis. Gels were made in-house and composed of 1% w:v agarose dissolved in TAE buffer (40mM Tris-Base, 20mM Acetic Acid, 1mM EDTA) supplemented with 0.02% Ethidium Bromide. These were cast and run in an Owl EasyCast B1A mini gel electrophoresis system (Thermo Fisher Scientific). Quick-Load Purple 1kb DNA ladder (New England Biolabs) was used for band reference. PCR or restriction digest products were mixed with 6x gel loading dye (New England Biolabs) and run at 100V for 30 minutes. Bands were visualized with UV light, excised, and purified using the Wizard SV Gel and PCR Clean-up System (Promega, A9281) as per manufacturer directions. Ligations were completed using the Quick Ligation Kit (New England Biolabs, M2200) according to manufacture protocol. Plasmids were transformed in NEB 5-alpha competent E. Coli following the NEB high efficiency transformation protocol (C2987H/C2987I). Transformed bacteria were grown on LB agar mixed with the appropriate antibiotic overnight. Monoclonal colonies were picked and expanded in LB broth for 12-18 hours, and then plasmid was purified using a QIAprep Spin Miniprep Kit (Qiagen, 27106) according to manufacture protocol.

Plasmids containing Kif19 cDNA were acquired from the RIKEN BioResource Center (73-76). Two missense point mutations present in the initial plasmid (c1411t and c2671t) were corrected using the Q5 Site-Directed Mutagenesis Kit (New England Biolabs, E0554S) according to manufacturer protocol. This processed Kif19 plasmid was utilized in PCR amplification with several primer pairs to generate multiple dsDNA Kif19 constructs: Full-length Kif19 (Kif19^FL^; forward primer: ATGAAGGACAGCGGGGACTCC; reverse primer: GTTATGCCGGGAGCATCCATCTTT), Kif19 lacking the C-terminal tail (Kif19^1-575^; forward primer: ATGAAGGACAGCGGGGACTCC; reverse primer: GTTATGCCGGGAGCATCCATCTTT), and Kif19 lacking the kinesin motor domain (Kif19^331-998^; forward primer: GAGGAGTCCCGGAACACC; reverse primer: GTTATGCCGGGAGCATCCATCTTT). These constructs were inserted into plasmids acquired from Addgene using restriction digest cloning: mClover (Clover2-N1, #54537) was digested with NheI and AgeI for Kif19 insertion, while tdTomato (tdTomato-C1, #54653) was digested with HindIII and KpnI. A plasmid expressing Kif19 attached to a C-terminal Myc tag was generated from the Clover2-N1 plasmid: we removed the Clover region by restriction digest with BsrGI and AgeI, and replaced it with a double-stranded DNA fragment encoding the Myc tag that purchased from Integrated DNA Technologies. pkMyc (Addgene, #19400) was used as a transfection control for these C-terminal Myc-tagged plasmids.

### Cell culture, siRNA treatment, and plasmid expression

DMS53, MDA-MB-231, U2OS, and HUVEC cells were all acquired from ATCC. DMS53 (CRL-2062) were cultured in Waymouth’s MB 742/1 Media (Thermo Fisher Scientific) supplemented with 10% fetal bovine serum (Atlanta Biologics) and 1% Penn-Strep (Thermo Fisher Scientific) at 37°C in the presence of 5% CO2. MDA-MB-231 (HTB-26) cells were cultured in DMEM (Thermo Fisher Scientific) supplemented with 10% fetal bovine serum, 1% Glutamax (Thermo Fisher Scientific, and 1% Penn-Strep at 37°C and 5% CO2. U2OS (HTB-96) cells were cultured in McCoy’s Modified 5A media (Thermo Fisher Scientific) supplemented with 10% fetal bovine serum, 1% Glutamax, and 1% Penn-Strep at 37°C and 5% CO2. HUVECs (PCS-100-010) were cultured in Medium 200 (Thermo Fisher Scientific) supplemented with 1% Gentamicin/Amphotericin B (Thermo Fisher Scientific) and 1x LVES (50x stock, Thermo Fisher Scientific) in flasks coated with ESGRO complete gelatin solution (Millipore Sigma) at 37°C and 5% CO2.

For passage of a 25cm^2^ flask, media was discarded and cells were washed with sterile PBS. Cells were detached using 1.5ml of 0.05% Trypsin (Thermo Fisher Scientific) incubated at 37°C for 5 minutes. Trypsin was neutralized with 8.5ml of media, and this solution was collected in a 10ml conical tube. If cells were being seeded for experiments, the cell concentration was estimated using 10µL of cell solution added to a Bright-Line 1492 hemocytometer (Hausser Scientific Partnership). Cells were passed into a new flask at a ratio of 1:3 for DMS53 and HUVECS, or 1:5 for U2OS and MDA-MB-231, and fresh media was added up to 10ml.

For *in vitro* Kif19 knockdown studies, cells were plated on a 24-well or 6-well plate at a density of 40k or 250k per well, respectively, 24 hours prior to transfection. For a 24-well plate, cells were transfected using 0.75ul Lipofectamine 3000 (Thermo Fisher Scientific) with 15 pmol Kif19 siRNA (Horizon Discovery, Kif19 SMARTpool of 4 siRNAs, L-004966-01), or a negative control siRNA (Sigma Aldrich, MISSION siRNA Universal Negative Control #1, SIC001). For a 6-well plate, 3.75 Lipofectamine 3000 and 50 pmol siRNA was used.

For plasmid transfection, the Lipofectamine 3000 kit (Thermo Fisher Scientific) was used with a modified protocol. Cells were plated on a 24-well plate at a density of 40k per well 24 hours prior to transfection. Each well was transfected using 50ul Opti-MEM (Thermo Fisher Scientific), 0.75ul Lipofectamine 3000, 1ul P3000 per 1ug plasmid, and ∼0.05pmol plasmid. The solution was added dropwise to each well.

### Western blotting

DMS53, MDA-MB-231, or U2OS cells were lysed 72 hours after transfection with siRNA or plasmids to generate whole cell lysates for western blot analysis. Cells were washed twice with 1x PBS and then lysed on ice with ice cold RIPA Buffer at a ratio of 100ul per 500k cells. After 20 minutes, the RIPA solution was collected in a 1.5ml microcentrifuge tube, and cell debris was spun down at 20,000xg for 10 minutes at 4°C using a 5810 R Centrifuge (Eppendorf). Protein concentration was estimated using the Pierce BCA Protein Assay Kit (Thermo Fisher Scientific, 23225) according to manufacturer protocol. The supernatant was aliquoted into fresh 1.5ml microcentrifuge tubes, flash frozen in Liquid N_2_, and stored at −80°C. At the time of use, lysate was mixed 3:1 v:v with 4x Laemmli sample buffer (Bio Rad) and 30:1 v:v with β-mercaptoethanol.

SDS-PAGE gels were made in-house using a Mini-Protean casting stand (Bio Rad). Gels were composed of a 10% acrylamide resolving layer underneath a 6% acrylamide stacking layer with combs for 10 or 15 wells. The resolving gel was composed of 4.0mL distilled water, 3.3mL 30% Acryl/Bis solution (VWR Life Science), 2.5mL 1.5M Tris Base pH 8.8, 0.1mL 10% Sodium Dodecyl Sulfate (SDS), 0.1mL fresh 10% Ammonium Persulfate, and 0.01mL TEMED. This solution was added to the cast, covered with a thin layer of pure ethanol, and allowed to polymerize for at least 30 minutes. Then the ethanol layer was poured off. The 6% acrylamide stacking gel was composed of 1.31mL distilled water, 0.4mL 30% Acryl/Bis solution (VWR Life Science), 0.25mL 1.0M Tris Base pH 6.8, 0.02mL 10% SDS, 0.02mL fresh 10% Ammonium Persulfate, and 0.002mL TEMED. This solution was added to the cast, a comb for 10 or 15 wells was inserted, and the solution was allowed to polymerize for at least 15 minutes.

SDS-PAGE gels were loaded into a Mini-protean Tetra Cell apparatus (Bio Rad) filled with in-house running buffer (25mM Tris-Base, 192mM Glycine, and 0.1% SDS in distilled water). Wells were loaded with 15ug of sample lysate, and 10ul of Precision Plus Protein Dual Color ladder (Bio Rad) was loaded into one well for reference. Gels were run for 90 minutes at 120V and 35mA. Gels were transferred onto a nitrocellulose membrane using a Mini Trans-Blot Cell system (Bio Rad) transfer system and an in-house transfer buffer (25mM Tris-Base, 192mM Glycine, and 20% Ethanol in distilled water). Transfer was run for 105 minutes at 100V and 476mA. All subsequent incubation steps were conducted on a shaker. Membranes were briefly incubated with Ponceau-S stain to visualize transferred lanes and trim membranes to necessary size. Membranes were then incubated in 5% w:v bovine milk in TBST buffer (150mM NaCl, 25mM Tris-HCl pH 7.5, and 0.05% Tween in distilled water) at room temperature (RT) for 1 hour. Subsequently, the 5% milk solution was discarded, and membranes were incubated with a solution containing primary antibodies diluted in 2% bovine milk in TBST at 4°C overnight. Membranes were then washed three times with TBST at RT for 10 minutes. A solution of secondary antibody in 5% milk in TBST was added at RT for 1 hour. Membranes were again washed three times with TBST at RT for 10 minutes. Membranes were developed with SuperSignal West Pico PLUS or Femto Chemiluminescent solution (Thermo Fisher Scientific) for 5 minutes, visualized on a ChemiDoc Touch Imaging System (Bio Rad), and analyzed using Image Lab software (Bio Rad).

### Cell viability assay

DMS53 are MDA-MB-231 cells were plated on a six-well dish and transfected with control or Kif19 siRNA as described above. 72 hours after transfection, cells were detached and collected as detailed above, with the following modifications: 300uL of Trypsin were used per well and 900uL of media were used to neutralize the trypsin. These cells were pelleted at 216xg for 5 minutes on a tabletop centrifuge and resuspended in 400µL of PBS. 50 µL of this solution was mixed with an equal volume of Trypan Blue (Thermo Fisher Scientific) and allowed to incubate for two minutes. Then, 10 µL of the solution was added to a Bright-Line 1492 hemocytometer (Hausser Scientific Partnership), and the number of stained (dead) and unstained (live) cells was recorded. The average of four cell counter fields-of-view was used to estimate the percent live and dead cells per condition

### Transwell invasion assay

On day 1, a 100µL aliquot of Matrigel Matrix (Corning Life Sciences, 354230) was moved from −20°C to 4°C to thaw overnight, and DMS53 or MDA-MB-231 cells were plated on a 6-well dish for transfection as detailed above. On day 2, cells were transfected with control or Kif19 siRNA. Four sterile 8um-pore membrane cell culture inserts (Cell Biolabs) were placed on a sterile 24-well dish in a cell culture hood. 50ul of an ice-cold 1:1 Matrigel:media solution was carefully added to the membrane of each insert under sterile conditions. The dish was stored at 37°C in the cell culture incubator overnight to allow the Matrigel solution to polymerize.

On day 3, cells were detached and collected as detailed above, with the following modifications: 300uL of Trypsin were used per well and 900uL of media were used to neutralize the trypsin. Cells were counted and 250k cells each of control and Kif19 siRNA-treated cells were collected in two new 15mL conical tubes. These cells were pelleted at 216xg for 5 minutes on a tabletop centrifuge. The supernatant was discarded, and cells were resuspended in 250ul of serum-free media. 100ul of cell solution (i.e. 100k cells) was carefully added to each of the Matrigel-coated membranes in the 24-well plate (two membranes per condition). Lastly, 400uL of media containing 10% FBS was added to the lower chamber of each well containing a membrane to act as a chemoattractant. The dish was then incubated at 37°C and 5% CO_2_.

Cells were fixed and stained on day 5 (72 hours following siRNA transfection) using a modified version of the protocol for the CytoSelect 24-Well Cell Invasion Assay, Colorimetric Format (Cell Biolabs, CBA-110). The Cell Stain Solution (Part No. 11002) was used to visualize the cells, and then brightfield images of randomized membrane sections were captured at 10x magnification using an EVOS Auto FL epifluorescent microscope (Life Technologies). FIJI image analysis software (77, 78) was used to determine the area covered by invading cells as a proportion of the total image area.

### Spheroid sprouting assay

DMS53 cells were transfected with a tdTomato transfection marker and control or Kif19 siRNA as described above. 48 hours after transfection, the cells were detached with trypsin and counted. To generate spheroids, cells were added to a 96-well PrimeSurface plate (S-BIO) at a concentration of 5k cells per well. 24-48 hours later, the spheroids were collected using a wide-orifice pipette tip and pooled in groups of five. Concurrently, HUVECs were cultured in a 24-well dish to 80-90% confluency. Each group of spheroids was added to a well containing HUVECs. After 24-48 hours of co-culture, the spheroids were imaged using an EVOS Auto FL epifluorescent microscope (Life Technologies) at 10x magnification. Spheroids were then scored based on presence or absence of tdTomato-positive cells sprouting from the spheroid edge.

### Fixation for immunofluorescence

DMS53, MDA-MB-231, or U2OS cells were grown on coverslips and fixed in an ice-cold or RT solution of BRB80 containing 4% paraformaldehyde (PFA), 0.1% Triton X-100, and 0.15% Glutaraldehyde for 15 minutes. Cells were briefly washed three times with PBS, and then incubated with an ice-cold or RT solution of 7.5mg/mL Sodium Borohydrite in PBS for 20 minutes to inactivate the glutaraldehyde. Cells were again washed three times with PBS, before being incubated with 0.5% Triton X-100 in PBS at RT for 5 minutes. Fixed cells were then incubated with blocking buffer (5% normal goat serum, 0.02% sodium azide, and 0.1% Triton X-100 in PBS) at RT for 1 hour. Primary antibodies were diluted in blocking buffer and incubated on the fixed cells at 4°C overnight. To remove excess primary antibody, cells were washed three times with PBST (0.05% Tween in PBS) at RT for 10 minutes. Cells were then incubated with Secondary antibodies diluted in blocking buffer at RT for 1 hour in the dark. Excess antibody was again removed by washing three times with PBST. Coverslips were mounted onto microscope slides using Immuno-Mount (Thermo Fisher Scientific) and allowed to dry overnight at 4°C.

### Confocal microscopy and general image pre-processing

Two confocal imaging systems were used in our assays. All E-cadherin and Vinculin assays, the microtubule assays in DMS53 cells, and all qualitative imaging of Kif19 plasmids were conducted on a 4-D spinning-disk confocal microscope (PerkinElmer) with 60x (1.4 NA) and 100x (1.4 NA) objectives and two excitation lines (488nm and 568nm) attached to a digital camera (Orca ER; Hamamatsu). The microtubule assays in MDA-MB-231 cells were conducted at the Einstein Analytical Imaging Facility on a SP5 AOBS (Leica) confocal microscope with a 63x (1.4 NA) objective, and three excitation lines (488nm, 543nm, and 633nm) attached to a point scanner (Inverted DMI6000; Leica).

To ensure images were quantitatively comparable, control and experimental conditions were imaged under identical conditions. Researchers were blinded to experimental conditions throughout imaging and subsequent analysis for immunofluorescent imaging assays. Images were processed and analyzed using FIJI iamge analysis software (77, 78). Initial preprocessing for all assays was done by first generating z-stack projections for each fluorescent channel using the maximum fluorescent intensities of each slice. Subsequently, we generated ROIs encircling each individual cell.

### Tubulin immunofluorescent analysis assays

72 hours prior to fixation, DMS53 or MDA-MB-231 cells were transfected with siRNA or Kif19 plasmids, as described above. For the low-dose nocodazole assay, DMS53 were additionally incubated with media containing 75nM nocodazole or DMSO control for two hours at 37°C prior to fixation. Cells were fixed following the protocol described above using solutions at RT.

For the tubulin tyrosination assays, DMS53 or MDA-MB-231 cells stained for α-tubulin and tyrosinated-tubulin were imaged and pre-processed as described above. For a given cell ROI, the total fluorescent intensity of each fluorescent channel was measured and corrected for background to generate the corrected total cell fluorescence (CTCF). The α-tubulin CTCF was used to represent the total microtubule polymer levels per cell. The tyrosinated-tubulin CTCF was divided by the α-tubulin CTCF to generate a ratio describing what proportion of the MT array was tyrosinated. This same process was completed for the tubulin detyrosination assays, except cells were stained for α-tubulin and detyrosinated-tubulin.

To examine a total microtubule polymer levels following nocodazole treatment or ectopic Kif19 expression, cells were stained for α-tubulin ± Myc (for Kif19 construct overexpression studies). Cells were imaged and preprocessed as described above, and the α-tubulin CTCF was calculated to represent the total microtubule polymer levels per cell.

### E-cadherin immunofluorescent analysis assays

72 hours prior to fixation, DMS53 or MDA-MB-231 cells were transfected with siRNA or Kif19 plasmids, as described above. Cells were fixed following the protocol described above using ice-cold solutions. To calculate whole cell E-cadherin, cells stained for E-cadherin and either actin (for Kif19 knockdown studies) or Myc (for Kif19 construct overexpression studies) were imaged and pre-processed as above, and the CTCF of the E-cadherin fluorescent channel was calculated. To calculate E-cadherin accumulation at cell-cell contacts, the FIJI brush tool was used to generate a 1um-wide ROI selection along the border of adjacent cells. The CTCF within this region was calculated and then corrected for the ROI area to account for variation in cell-cell contact area.

### Focal adhesion analysis

72 hours prior to fixation, DMS53 or MDA-MB-231 cells were transfected with siRNA or Kif19 plasmids, as described above. Cells were fixed following the protocol described above using ice-cold solutions. Cells stained for Vinculin and either actin (for Kif19 knockdown studies) or Myc (for Kif19 construct overexpression studies) were imaged and pre-processed as in the previous section. The area of each cell ROI was measured and stored. We then used a series of FIJI processes to generate a mask displaying only the Vinculin focal adhesions. To enhance the visibility of vinculin adhesions, we ran “enhance contrast” with saturated pixels at 0.35%, followed by “subtract background” with a rolling ball radius of 50 pixels, then “despeckle,” and lastly “8-bit.” To create masks from these processed images, the FIJI “auto local threshold” function was run with the Phansalkar method and a radius at 20. To generate a list of ROIs for each focal adhesion in the mask, the FIJI “analyze particles” function was run with a size range of 0.04-20um^2^ (for DMS53) or 0.12-40um^2^ (for MDA-MB-231) and circularity range of 0.00-1.00. Subsequently, this list of ROIs was applied over the original Vinculin z-projection and the ROIs were manually edited to ensure they accurately represented the foal adhesions on the image. The size measurements for each individual focal adhesion ROI was saved. To calculate focal adhesion density, the sum of all focal adhesion ROIs within a cell was divided by the total cell area. To calculate effective adhesion density, the ImageJ brush tool was used to generate a 2um border along the edges of cells, excluding areas of cell-cell contact. The sum of all focal adhesion ROIs within a cell was divided by this effective cell area.

### Statistical analysis

All statistical analyses were completed in GraphPad Prism 8.X (GraphPad Software). Differences between treatments were analyzed using a student’s t-test to compare two groups. Outliers were removed from independent experiment data sets using the ROUT method with Q=1%. To account for environmental variation between experiments, data from each experiment was normalized to the mean value of that experiment’s control condition. Means were determined to be significant if p<0.05.

## Results and discussion

### Kif19 depletion reduces cell invasiveness *in vitro*

Kif18A (ENSG00000121621) and Kif18B (ENSG00000186185) regulate chromosomal segregation during mitosis and promote oncogenesis chiefly by increasing cell survival and proliferation (57, 59, 61). Our preliminary studies indicate that Kif19 (ENSG00000196169) expression correlates with decreased overall survival probability in several cancers (**Fig S1**), so we initially sought to determine if Kif19 regulates cell viability in cancer cells. We reviewed several databases for cell lines with high Kif19 expression and selected two cell lines: DMS53 and MDA-MB-231. These cell lines were chosen for their high Kif19 expression, adherent morphology, and motility. DMS53 cells are a human primary small cell lung cancer line, while MDA-MB-231 cells are derived from a human metastatic breast adenocarcinoma. We optimized Kif19 knockdown in the cell lines using a pool of fours siRNAs (Horizon Discovery, L-004966-01); verification by western blot indicated approximately 61.4% and 41.9% knockdown in DMS53 and MDA-MB-231 cells, respectively (**Fig S2A,B**). Following Kif19 knockdown, we observed no change in the viability of MDA-MB-231 or DMS53 cells (**Fig 1A**). This may reflect the fact that, unlike other kinesin-8 members, Kif19 is not associated with mitosis (64, 67-71). Kif19 is reported to localize with MTs at the tips of cilia, but neither DMS53 nor MDA-MB-231 cells are ciliated (64). It was previously reported that a Kif18A mutant lacking a functional nuclear localization sequence (NLS) localized to the cell periphery, where it suppressed MT growth and dynamics (79). We theorized that the Kif19 expressed in our cell lines could similarly be localizing to and acting upon MTs in the cell periphery. As MT dynamics at the cell edge can extensively regulate cell movement, we sought to determine if Kif19 regulates cell invasion (31, 38, 80). Additional analysis of Kif19 expression in publicly available cancer patient datasets indicated that, in colorectal and breast carcinoma, Kif19 expression was significantly higher in samples from metastases than from primary tumors (**Fig S3A,B**). DMS53 and MDA-MB-231 cells treated with Kif19 or control siRNA were used in a Matrigel transwell invasion assay. Kif19 knockdown led to a 45% (p<0.0001) and 58% (p<0.0001) decrease in invading cell area for DMS53 and MDA-MB-231 cells, respectively (**Fig 1C**). To characterize Kif19’s effect on invasion in a more physiologically relevant system, we seeded siRNA-treated DMS53 spheroids onto a monolayer of primary human umbilical vein endothelial cells (HUVECs) (**Fig 1D**). Spheroids treated with control siRNA often exhibited “sprouting” cells that invaded into the surrounding HUVEC monolayer. However, Kif19 depletion led to a significant decrease in the number of spheroids exhibiting sprouting and invasion (79.3% vs 45.1% spheroids sprouting for control vs Kif19 siRNA, respectively, p=0.017) (**Fig 1E**). From these data, we propose that the pro-carcinogenic effects of Kif19 are mediated by regulation of cell invasiveness, rather than regulation of the cell cycle as seen with other kinesin-8 members.

**Fig 1.**
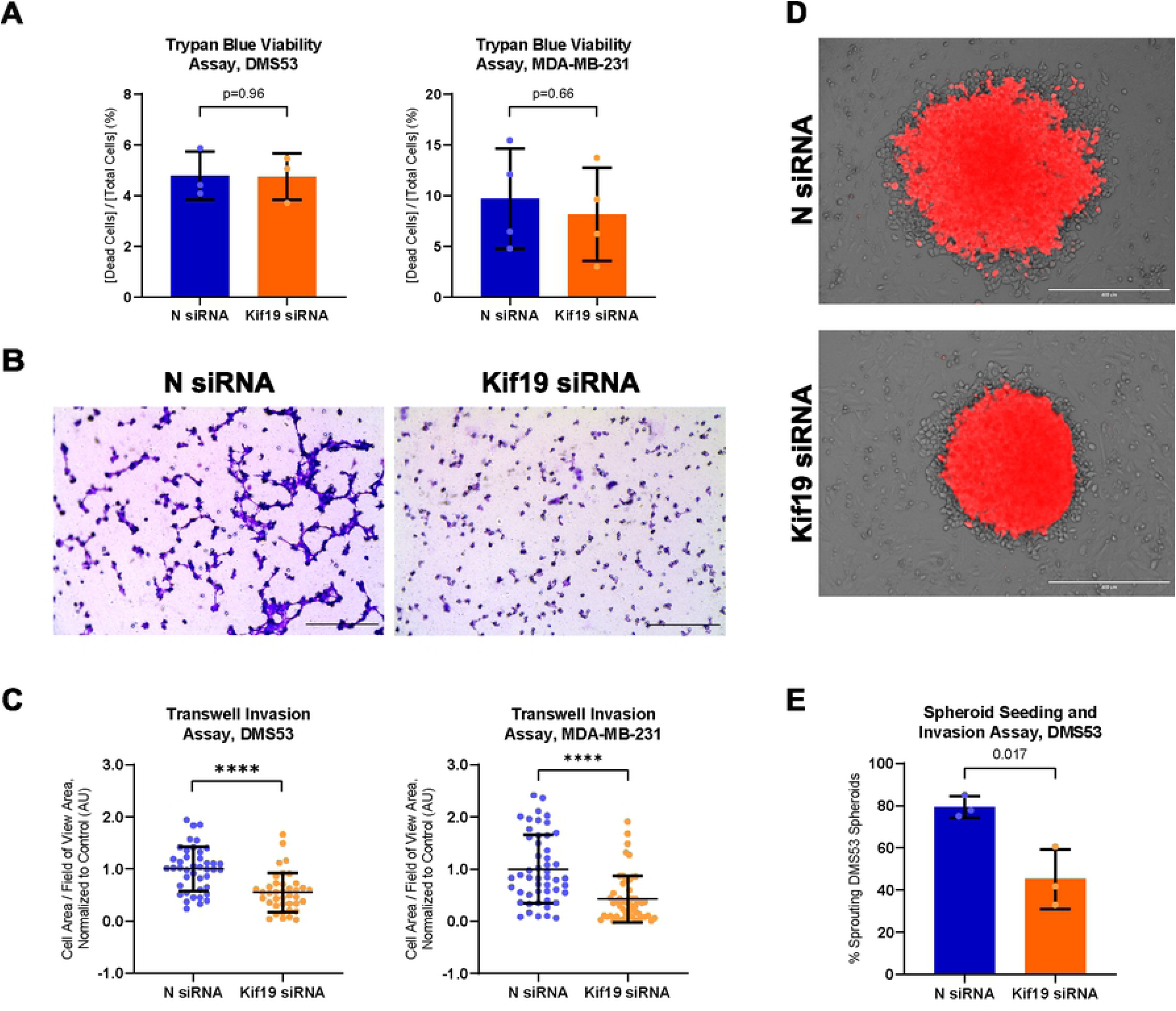
Kif19 regulates cell invasion but not viability. **A)** Graphs describing the viability of DMS53 cells (left) and MDA-MB-231 cells (right) transfected with control (N) or Kif19 siRNA as analyzed by Trypan Blue staining. Graphs show the results of three and four independent experiments for DMS53 and MDA-MB-231, respectively, and indicate that there is no significant change in viability following Kif19 knockdown. **B)** Representative images of fixed invading N and Kif19 siRNA-treated DMS53 cells from transwell Matrigel invasion assays. Scale bars represent 200um **C)** Graphs depicting the percent cell area covered by invading DMS53 (left) and MDA-MB-231 (right) cells transfected with N or Kif19 siRNA in transwell Matrigel invasion assays. Five independent repetitions were completed per cell line (n=∼9 randomly selected fields of view each). Data indicates that Kif19 siRNA treatment significantly decreases the area covered by invading cells in DMS53 and MDA-MB-231 cells. **D)** Representative brightfield images of DMS53 cells transfected with siRNA and tdTomato (red) seeded on a HuVEC monolayer and exhibiting sprouting (top, N siRNA) or no sprouting (bottom, Kif19 siRNA). Scale bars represent 400um. **E)** Graph depicting the percentage of DMS53 spheroids seeded on a HuVEC monolayer that exhibited sprouting within 5 days post-seeding, which indicates that Kif19 siRNA treatment significantly decreases the number of sprouting spheroids. Data represent the sum of three independent repetitions (n=∼25 spheroids each). Graph error bars depict mean ± standard deviation; p-values calculated with a Student’s t-test; ****, p<0.0001.

### Kif19 localizes to the nucleus and peripheral MTs

To understand how Kif19 alters cell invasion, we examined the protein’s localization within the cell. While several commercial Kif19 antibodies are available, these proved inconsistent for IF imaging. Two, AV34084 and HPA073303, bound exogenous, plasmid-expressed Kif19 on western blot, but not on IF imaging (**Fig S2C,D**). We thus focused on examining the localization of plasmid-based Kif19 constructs. These studies were completed in DMS53 cells as well as U2OS cells, which are an osteosarcoma cell line used for their low endogenous Kif19 expression and well-defined MT array. In initial live imaging studies, fluorescent full-length Kif19 constructs (Kif19^FL^) localized to both the nucleus and MTs (**Fig 2A**). We then examined localization of Kif19^FL^ along MTs using IF imaging. We observed a gradient of Kif19 concentration along MTs, with accumulation at MT plus-ends in the cell periphery (**Fig 2B**). This is characteristic of kinesin-8 members and is often reliant on the kinesin’s CTT as well as the motor domain (52-54). As expected, neither a Kif19 construct lacking the CTT (Kif19^1-575^) nor a construct lacking the kinesin motor domain (Kif19^331-998^) concentrated along microtubules, suggesting that Kif19 behaves similar to other kinesin-8s (**Fig 3**). This accumulation at MT ends is often due to the presence of a secondary MT-binding site within the kinesin-8 CTT, which results in highly processive movement of kinesin-8s along microtubules (50, 53, 81). We observed that Kif19^331-998^ chiefly localized to the nucleus, but was present on microtubules when highly overexpressed, suggesting that a secondary binding site may be present (**Fig 3C**).

**Fig 2.**
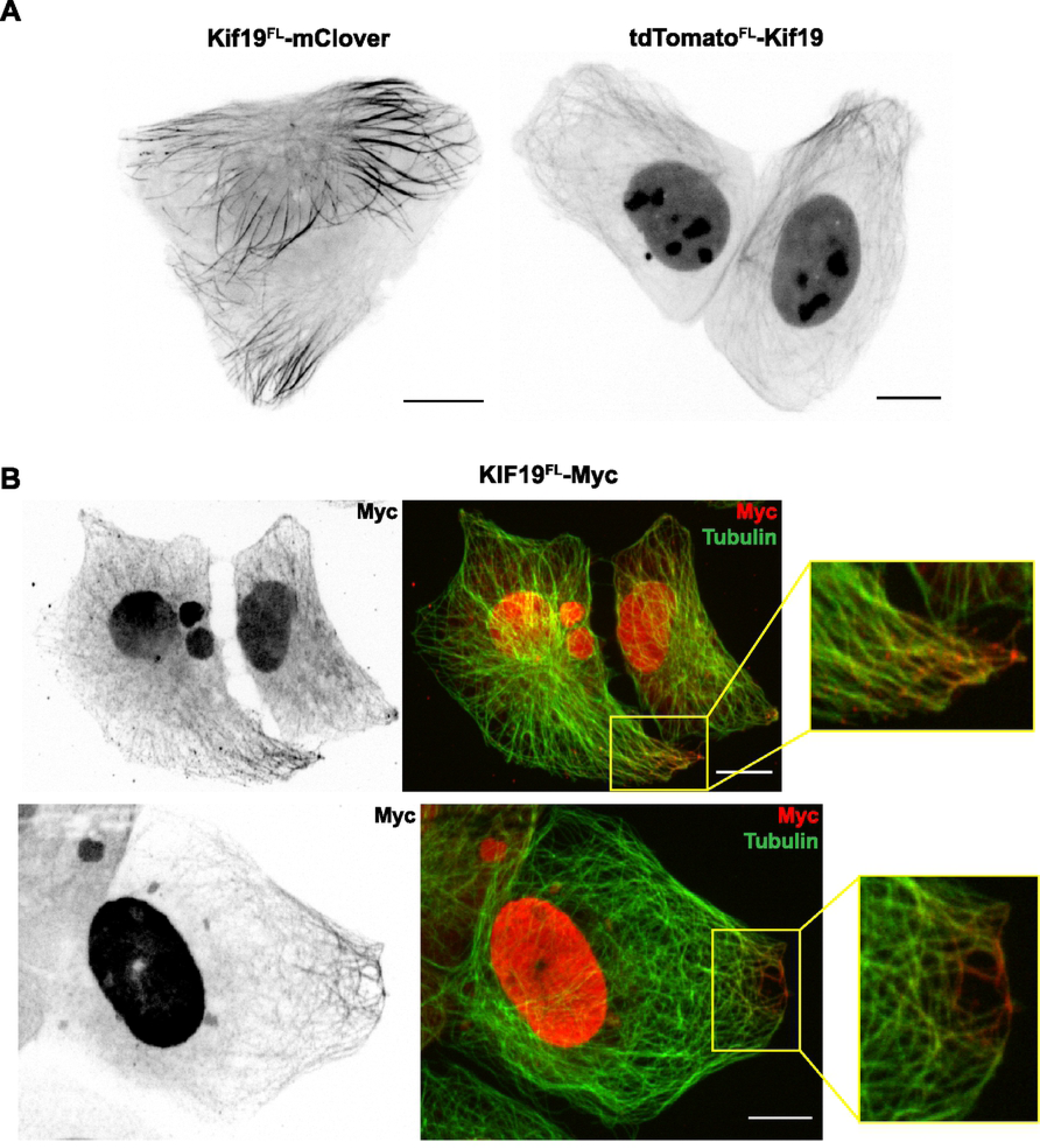
Kif19 localized to the nucleus and peripheral MTs. **A)** Confocal imaging of live U2OS cells expressing Kif19^FL^-mClover (left) and tdTomato-Kif19^FL^ (right) exhibiting colocalization with the nucleus and peripheral MTs. **B)** Confocal imaging of a U2OS cells expressing Kif19^FL^-Myc that were fixed and stained for Myc and α-tubulin. Insets highlight colocalization of ectopic Kif19 with peripheral MTs. Scale bars represent 10um.

**Fig 3.**
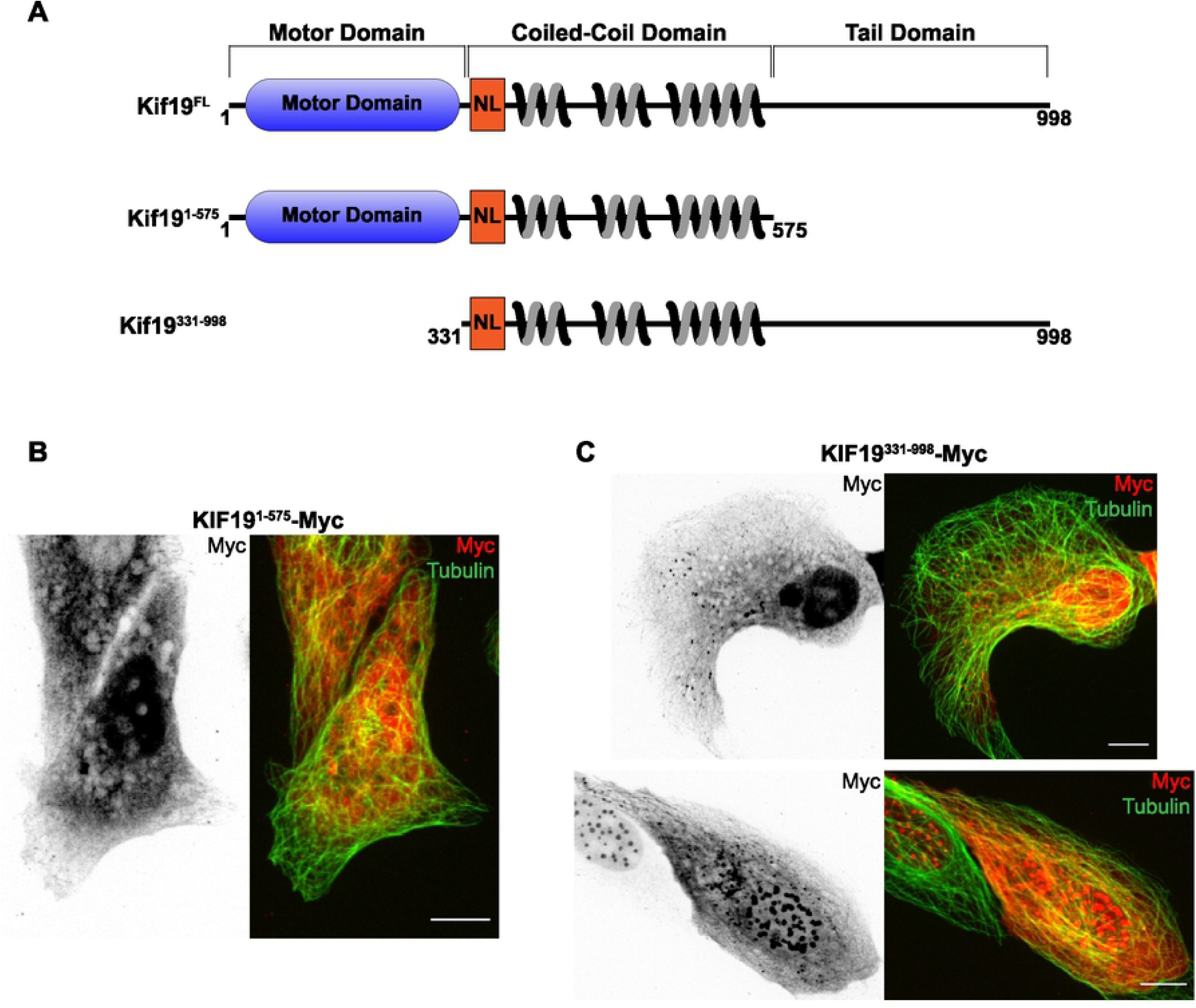
Kif19 localization along MTs relies on the full-length protein. **A)** Diagram illustrating the structural components of the Kif19 monomer, as well as the truncated constructs used in this report (NL: neck linker). **B)** Confocal imaging of a DMS53 cell expressing Kif19^1-575^-Myc that was fixed and stained for Myc and α-tubulin. Note loss of MT colocalization. **C)** Confocal images of DMS53 cells transfected with Kif19^331-998^-Myc that were fixed and stained for Myc and α-tubulin. Note nuclear localization (top) and MT colocalization (bottom). Scale bars represent 10um.

### Kif19 has MT stabilizing and destabilizing properties

The highly processive nature of kinesin-8 proteins allows for an activity termed “length-dependent depolymerization,” in which longer MTs accumulate a greater number of kinesin-8 proteins at the MT plus-end and are thus more likely to undergo depolymerization (82, 83). Kif19’s motor domain was previously reported to rapidly depolymerize purified GMPCPP-stabilized MTs, and we hypothesized that overexpression of Kif19 constructs in cells would similarly depolymerize MTs (64, 65). However, DMS53 and U2OS cells expressing Kif19^FL^ or Kif19^1-575^ often contained a complete MT array and only rarely exhibited apparent MT depolymerization (**Fig 2 and Fig 4A,B**). We then quantified how Kif19 plasmids (described in Methods) affected the MT array by fixing DMS53 cells expressing Myc-tagged Kif19 constructs and staining them for Myc and tubulin. IF analysis indicated that overexpression of Kif19^FL^ resulted in a 18.5% decrease in total polymerized MTs compared to vector control (p<0.0001) (**Fig 4C**). Kif19^FL^ had a significantly greater impact on the MT array than Kif19^1-575^, even though both were observed to depolymerize MTs (18.5% vs 6.3% decrease compared to control for Kif19^FL^ vs Kif19^1-575^, respectively, p=0.0065). This suggests that the Kif19 CTT is important for the protein’s *in vivo* MT-depolymerizing activity, in addition to its localization.

**Fig 4.**
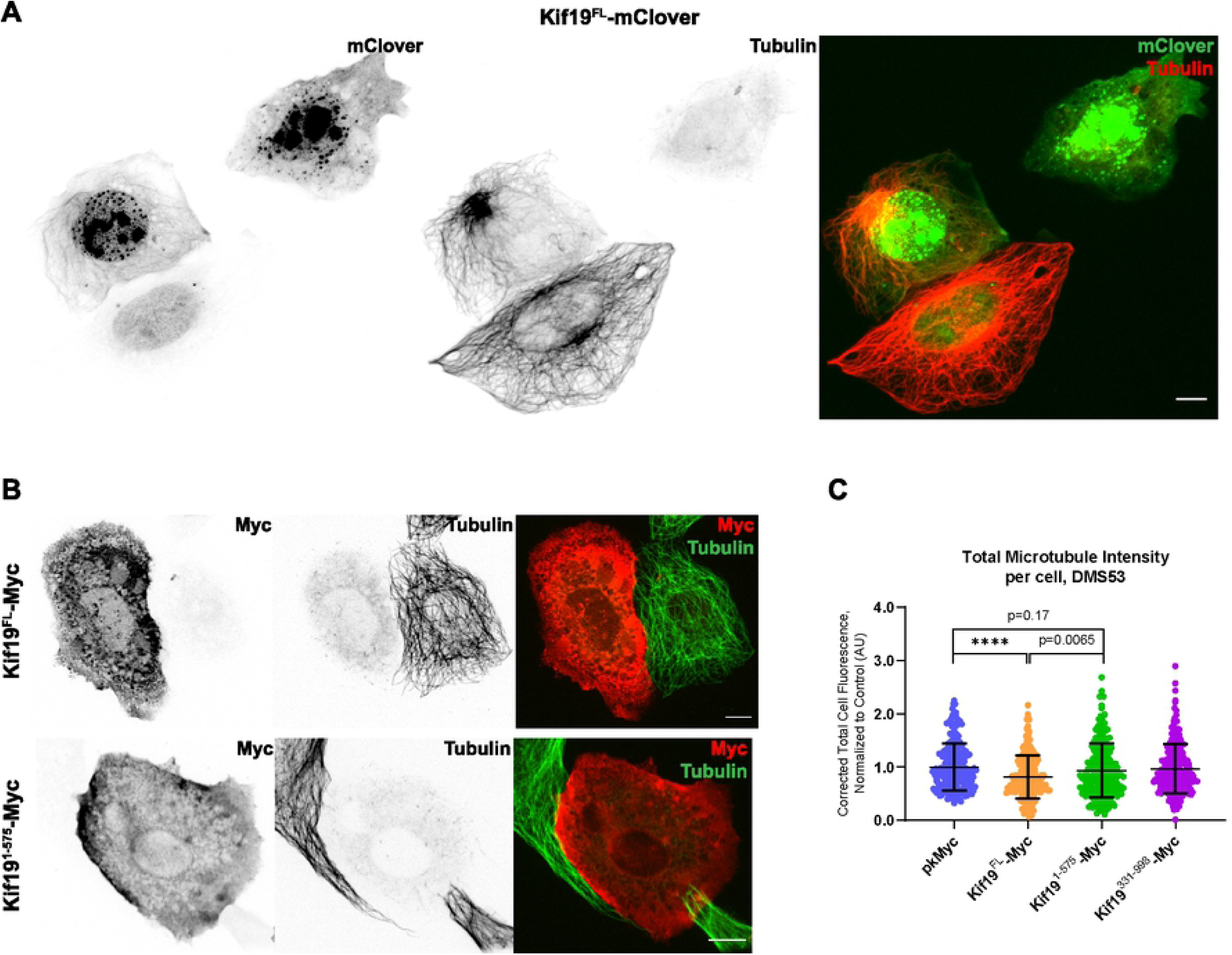
Kif19 has MT depolymerizing activity. **A)** Epifluorescent imaging of U2OS cells expressing Kif19^FL^-mClover that were fixed and stained for α-tubulin, which illustrates how the level of ectopic Kif19 expression can alter the MT array. **B)** Confocal images of DMS53 cells expressing Kif19^FL^-Myc (top) or Kif19^1-575^-Myc (bottom), which were fixed and stained for Myc and α-tubulin, displaying destruction of MT array. **C)** Quantitative image analysis of individual DMS53 cells expressing Myc-tagged constructs indicates that cells expressing Kif19^FL^-Myc exhibit a significant decrease in total polymerized MTs compared to cells expressing pkMyc control or Kif19^1-575^-Myc. Data were collected from 7 independent experiments (n=∼30 cells each). Graph error bars depict mean ± standard deviation; p-values calculated with Student’s t-test; ****, p<0.0001. Scale bars represent 10um.

Given our observed MT-depolymerization by Kif19, we theorized that Kif19 knockdown would increase the levels of polymerized MTs. Oddly, we observed that Kif19 depletion by siRNA in DMS53 and MDA-MB-231 cells had no significant effect on total polymerized MT levels (**Fig 5A**). However, this result is not outside the realm of possibility for Kif19. In addition to MT depolymerization, kinesin-8 members have been shown to stabilize microtubules, promote MT rescue, and decrease MT shrinkage velocity (49, 50, 84, 85). To determine whether Kif19 alters MT dynamics, we examined how the modification status of the tubulin array changes with Kif19 expression. IF analysis was performed on siRNA-treated cells fixed and stained for total tubulin and either tyrosinated-tubulin or detyrosinated-tubulin. The tyrosination status of tubulin is commonly used as a marker for MT stability; tyrosinated MTs are considered more dynamic, while detyrosinated MTs are considered more stable (86). By calculating the ratio of modified-to-total tubulin, one can assess how a treatment alters the dynamicity of the MT array. We observed that Kif19 depletion in DMS53 cells produced an 18.4% increase in the proportion of tyrosinated-tubulin (p<0.0001) and a corresponding 21.4% decrease in detyrosinated-tubulin (p<0.0001) (**Fig 5B,C**). Similarly, Kif19 knockdown in MDA-MB-231 cells resulted in a 9.3% increase in tyrosinated-tubulin (p<0.0001) and a corresponding 6.8% decrease in detyrosinated-tubulin (p=0.0021) (**Fig 5B,C**). This implies that Kif19 is targeting dynamic (i.e. tyrosinated) MTs, either by selective depolymerization or stabilization.

**Fig 5.**
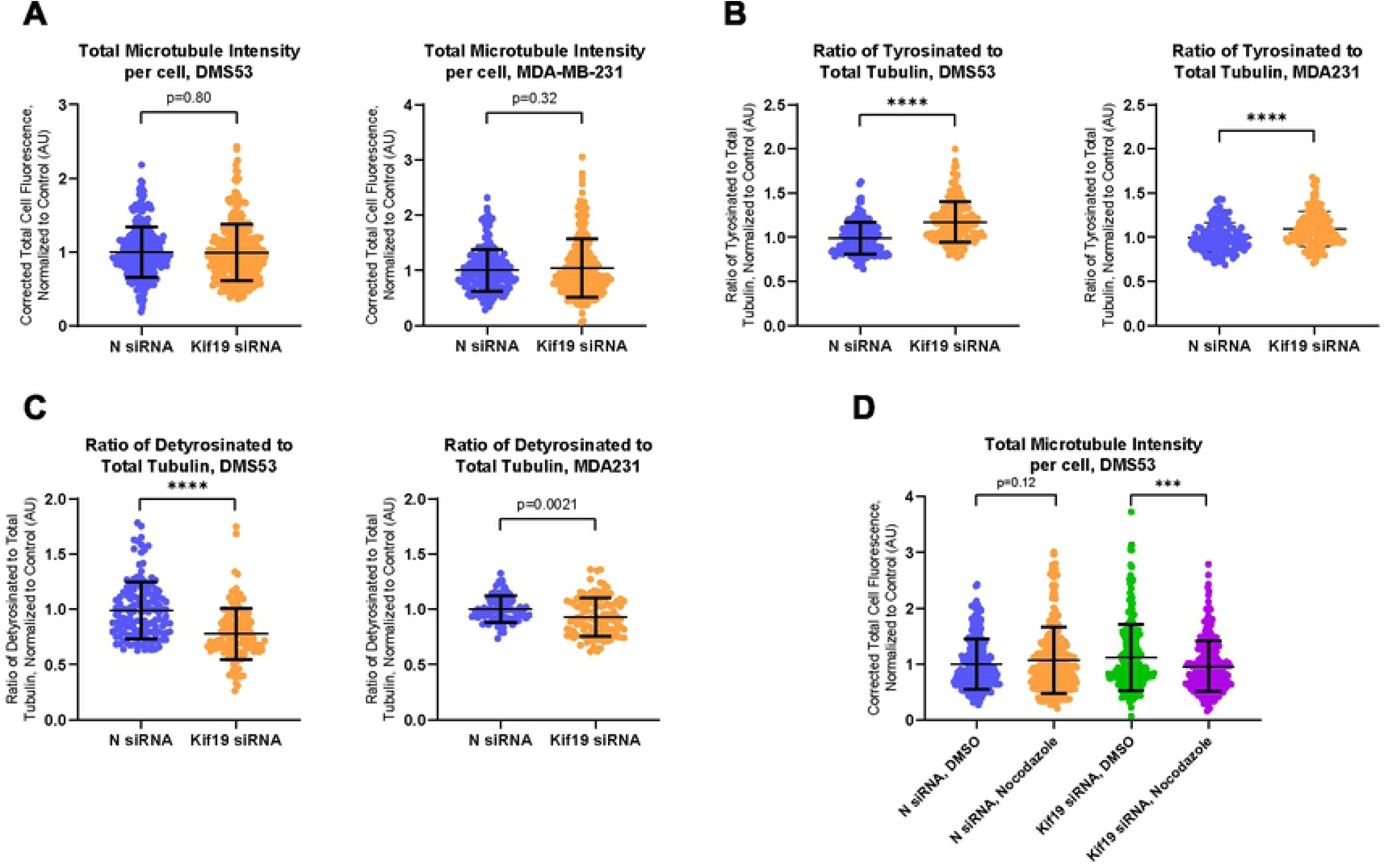
Kif19 exhibits MT-stabilizing activity. **A)** Quantitative image analysis of DMS53 (left) and MDA-MB-231 (right) cells indicates that Kif19 knockdown does not significantly alter total polymerized MT levels compared to control. Data were collected from seven experiments per cell line (n=∼40 cells each). **B)** Quantitative image analysis of DMS53 (left) and MDA-MB-231 (right) cells reveals that there is a significant increase in the ratio of tyrosinated:total tubulin following treatment with Kif19 siRNA. Data were collected from four experiments per cell line (n=∼40 cells each). **C)** Quantitative image analysis of DMS53 (left) and MDA-MB-231 (right) cells reveals that there is a significant decrease in the ratio of detyrosinated:total tubulin following treatment with Kif19 siRNA. Data were collected from three experiments per cell line (n=∼40 cells each). **D)** Quantitative image analysis of individual DMS53 cells transfected with N or Kif19 siRNA and treated with 75nM nocodazole or DMSO control. Data indicate that nocodazole treatment decreased total polymerized microtubules in Kif19 siRNA-treated cells, but not in N siRNA-treated cells. Data were collected from three independent experiments (n=∼80 cells each). Graph error bars depict mean ± standard deviation; p-values determined by Student’s t-test; ***, p<0.001; ****, p<0.0001.

To test whether Kif19 directly affects the stability of MTs, siRNA-treated DMS53 cells were subjected to a low dose of nocodazole or DMSO control for two hours prior to fixation and staining for tubulin. This low concentration of nocodazole inhibits MT dynamics without significantly depolymerizing the MT array (87-89). As expected, control siRNA-treated cells exhibited no significant difference in the amount of polymerized MTs between the DMSO and nocodazole treatments (7.2% change, p=0.12) (**Fig 5D**). However, DMS53 cells depleted of Kif19 showed a slight, but significant, decrease in polymerized MTs following nocodazole treatment, implying that the MT array is more labile (14.4% change, p=0.0004). Taken together, these data imply that Kif19 has a mixed effect on MTs, with both stabilizing and depolymerizing capabilities, similar to other kinesin-8s. We theorize that this dual function could allow Kif19 to maintain proper MT length in cilia without completely disrupting the array. However, in non-ciliated cancerous cells, aberrant Kif19 expression may disrupt MT dynamics in the cell periphery.

### Kif19 regulates focal adhesion size

MT dynamics are essential to FA regulation. MTs are guided towards adhesion sites by growth along actin stress fibers, and MTs undergo frequent, repeated targeting events upon reaching FAs (90, 91). Notably, these dynamic MT cycles appear to be required for FA turnover, as either pharmacological stabilization or depolymerization of MTs inhibits FA turnover (92). Given Kif19’s localization to MTs, impact on cell motility, and effect on MT dynamics, we sought to determine if Kif19 modifies FAs. We treated cells with control or Kif19 siRNA before fixing them and staining them for Vinculin (ENSG00000035403), a ubiquitous FA component. Using quantitative IF analysis, we determined that Kif19 depletion by siRNA in DMS53 cells led to a 19.5% increase in FA density (total FA area / total cell area) (p=0.013) (**Fig 6B**). This appears to be primarily mediated by an increase in the average FA size per cell (12.3% increase, p=0.0015), with only a slight, non-significant increase in the number of adhesions per cell (4.5% increase, p=0.40) (**Fig 6E**). Similarly, Kif19 depletion in MDA-MB-231 cells led to a 13.5% increase in FA density (p=0.0012), though individual FA size and number could not be accurately calculated due to extensive overlapping between adjacent adhesions (**Fig 6C and Fig S4A**). Overexpression of myc-tagged Kif19^FL^ in DMS53 cells led to a 16.3% decrease in FA density (p=0.0060), supporting our knockdown data (**Fig 6D**). However, this change in density was meditated by decreases in both FA number (6.3% decrease, p=0.44) and average FA size (5.9% decrease, p=0.11), with neither variable significant on its own (**Fig 6F**). Notably, overexpression of myc-tagged Kif19^1-575^ or Kif19^331-998^ did not significantly alter FA density compared to control, implying that the full-length protein is necessary for the observed effects (6.4% decrease and 1.2% increase in FA density for Kif19^1-575^ and Kif19^331-998^, respectively, p=0.29 and p=0.85).

**Fig 6.**
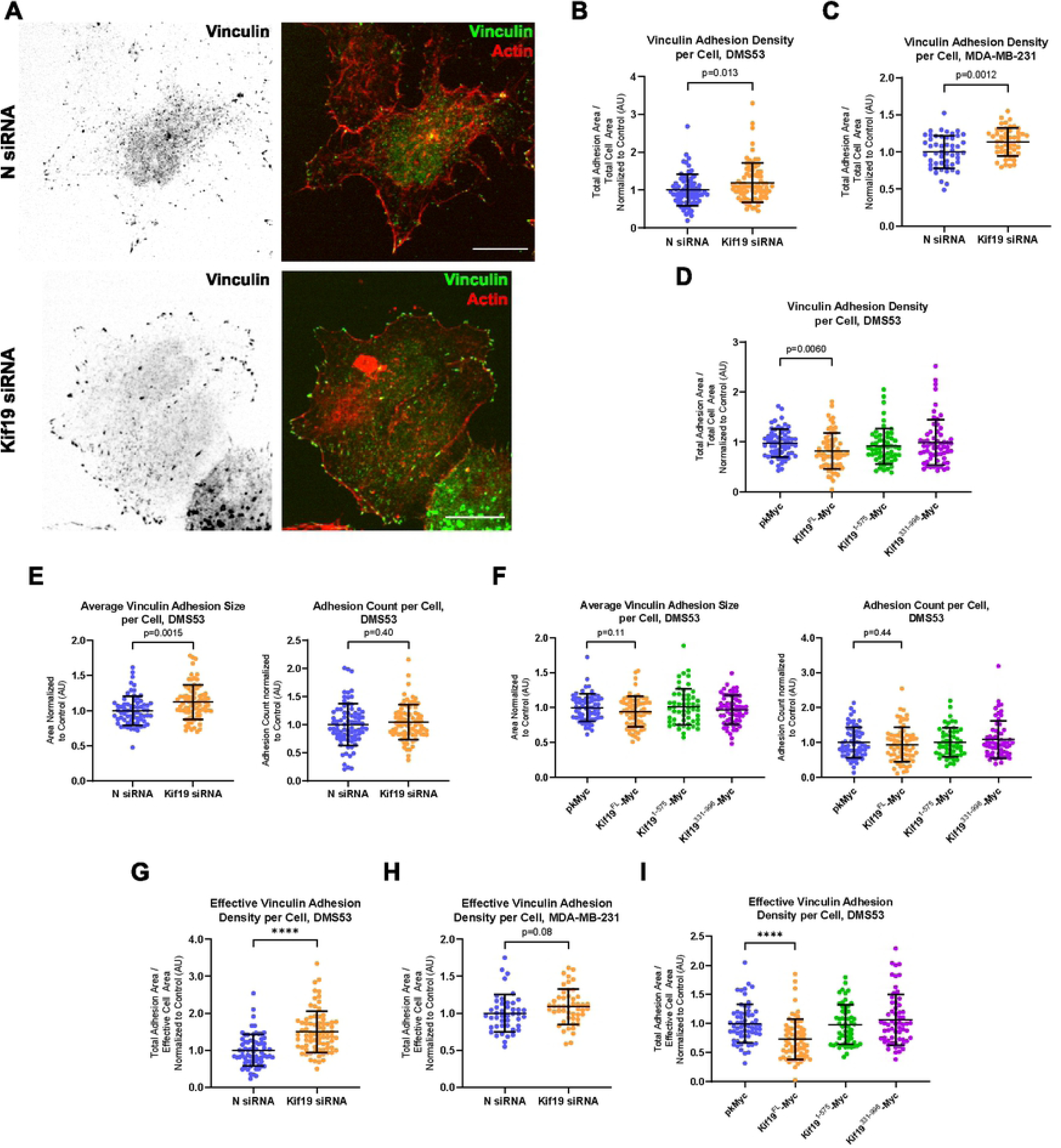
Kif19 regulates FA density. **A)** Representative confocal imaging of DMS53 cells treated with N or Kif19 siRNA, fixed, and stained for Vinculin and Actin. Scale bars represent 10um. **B-C)** Quantitative image analysis of individual DMS53 **(B)** and MDA-MB-231 **(C)** cells reveals that cells treated with Kif19 siRNA have significantly greater FA density compared to cells treated with N siRNA. Data were collected from four and three independent experiments (n=∼20 cells each) for DMS53 and MDA-MB-231, respectively. **D)** Quantitative image analysis of individual DMS53 cells expressing Myc-tagged constructs shows a significant decrease in the FA density of cells expressing Kif19^FL^-Myc compared to pkMyc control. **E)** Additional analyses of the data collected from DMS53 cells treated with N or Kif19 siRNA displayed in B indicates that Kif19 siRNA-treated cells have a significant increase in the average FA size per cell, but not in FA number per cell, compared to control. **F)** Additional analyses of the data collected from DMS53 cells expressing Myc-tagged Kif19 constructs displayed in D indicates that there is no significant change in the average FA size or FA number per cell between cells expressing Kif19^FL^-Myc and pkMyc control. **G-I)** Additional analyses of the data displayed in graphs B-D with FA density corrected for “effective” area rather than whole cell area (see **Fig S4**). Data suggests that Kif19 siRNA treatment significantly increases effective FA density in DMS53 **(G)**, but not MDA-MB-231 **(H)**, cells compared to N siRNA treatment. DMS53 cells expressing Kif19^FL^-Myc have significantly decreased effective FA density compared to pkMyc control **(I)**. FA density defined as [total FA area per cell]/[total cell area]. Graph error bars depict mean ± standard deviation; p-values determined by Student’s t-test; ****, p<0.0001.

DMS53 cells have a distinct FA distribution: FAs are almost exclusively at the cell edge and are excluded from sites of cell-cell contacts (**Fig 6A**). We qualitatively observed that Kif19 knockdown in DMS53 cells increased cell-cell contact area and thus decreased the area over which FAs were observed (**Fig S4B**). To correct for this, we calculated an “effective” cell area consisting of a 2um band along the cell perimeter, excluding areas of cell-cell contact (**Fig S4C**). The increase in FA density following Kif19 knockdown was over 2-fold greater within this effective area compared to the whole cell area (49.9% increase in effective FA density compared to control, p<0.0001) (**Fig 6G**). A similar effect was observed with overexpression of Kif19^FL^ plasmids in DMS53 (26.7% decrease in effective FA density compared to control, p<0.0001) (**Fig 6I**). However, there was no difference after correcting for effective area in MDA-MB-231 cells after Kif19 knockdown (8.9% increase in effective FA density compared to control, p=0.083) (**Fig 6H**). This may be because FAs in MDA-MB-231 cells are found throughout the cytoplasm and do not appear to be affected by cell-cell contacts, unlike DMS53. Overall, these data support our theory that Kif19’s effect on peripheral MTs is related to its impact on cell motility. FA size and stability are relevant factors in the invasiveness of cancer cells, with smaller, dynamic adhesions being associated with greater aggression and invasiveness (93, 94).

Multiple mechanisms have been implicated in MT regulation of FA turnover. As FAs are tension-dependent structures, it has been proposed that MTs may relax the adhesion. This could be due to localized alterations in Rho GTPase activity by MTs, which is potentially mediated by KANK1/2 (95). The KANK proteins facilitate transient MT capture at FAs by cross-linking MTs with the FA component Talin (95). Disruption of this cross-linking leads to excessive RhoA-mediated myosin II activation and subsequent FA stabilization (96). Another possibility is the delivery of a relaxing factor by MTs. APC is a large protein transported along MTs to FAs that was recently shown to be important for proper FA disassembly and may be a factor involved with FA relaxation (97, 98). Beyond FA relaxation, MTs have been implicated in degrading the ECM around FAs by delivering exocytic particles containing matrix metalloproteinases (MMPS). This is mediated by the CLASP proteins, which localize to the plasma membrane surrounding FAs and transiently capture MTs, leading to the delivery and exocytosis of MMPs (99). We propose that Kif19’s alteration of MT dynamics at the cell edge changes FA assembly and disassembly, likely by modifying tension at FAs, resulting in more dynamic adhesions and allowing for greater cell motility.

### Kif19 regulates E-cadherin concentration at cell-cell contacts

We observed a qualitative increase in cell-cell contacts within areas of moderate confluency in Kif19-depleted DMS53 cells relative to controls (**Fig S4B**). Cell-cell AJs are an important factor in cancer progression that can be regulated by both MTs and FAs (32, 100), so we investigated the possible regulation of cell-cell adhesion by Kif19. IF analysis of siRNA-treated DMS53 cells fixed and stained for the AJ component E-cadherin (ENSG00000039068) revealed that Kif19 knockdown resulted in significantly more E-cadherin within the whole cell (17.6% increase, p=0.0037) and at cell-cell contacts (43.0% increase, p<0.0001) (**Fig 7B,E**). A similar phenotype was observed following Kif19 depletion from MDA-MB-231 cells (26.3% and 19.5% increase in E-cadherin for whole cell and cell-cell contacts, respectively, p<0.0001 and p=0.0050) (**Fig 7C,F**). Overexpression of full length Kif19 in DMS53 cells led to lower E-cadherin levels at cell-cell contacts, but not whole cell expression (5.8% and 17.7% decrease in E-cadherin for whole cell and cell-cell contacts, respectively, p=0.40 and p=0.0082) (**Fig 7D,G**). Notably, while overexpression of truncated Kif19 plasmids did not significantly alter E-cadherin levels at cell-cell contacts, Kif19^331-998^ overexpression did significantly decrease whole cell E-cadherin (16% decrease, p=0.021). The underlying mechanism of this decrease remains unclear.

**Fig 7.**
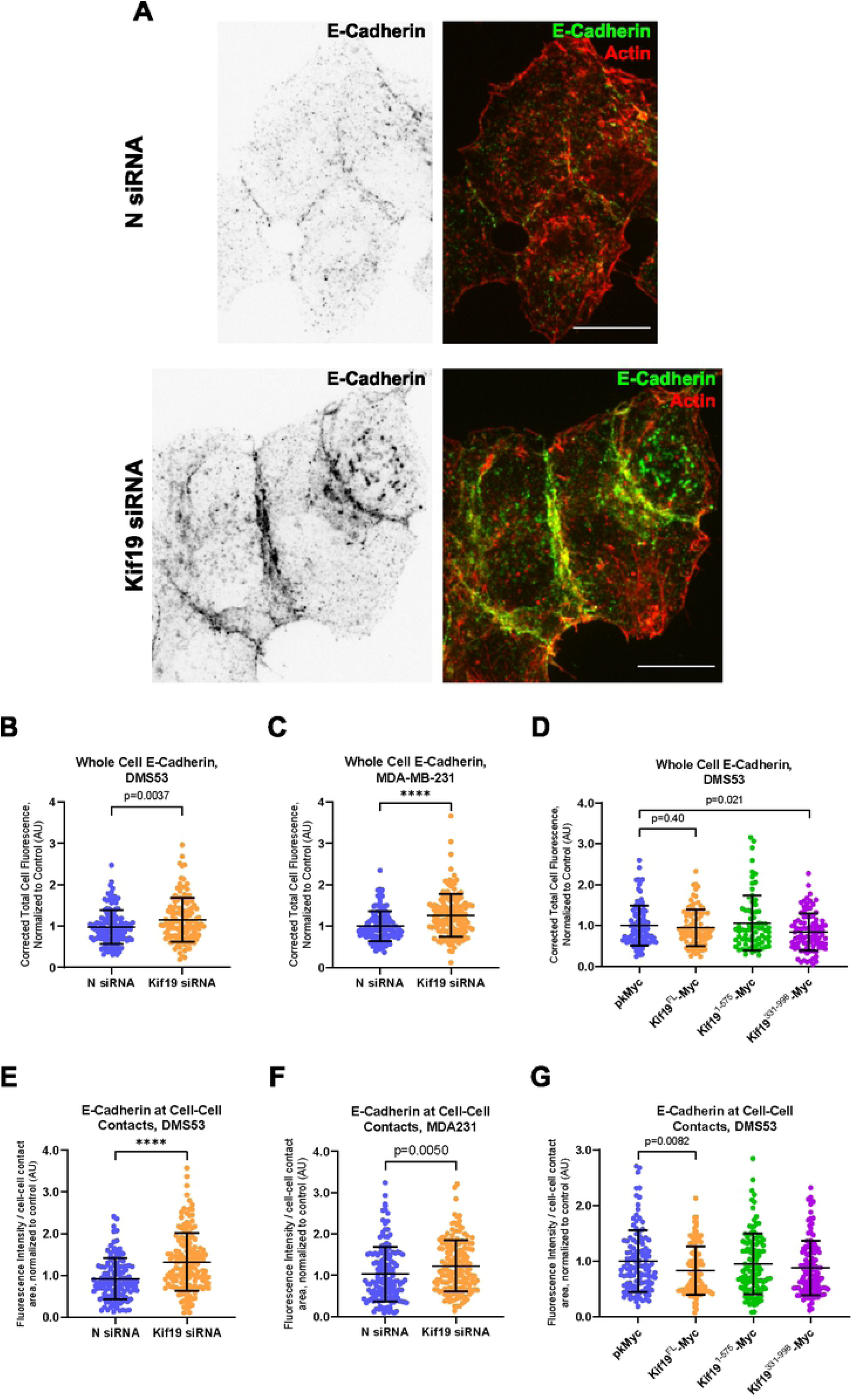
Kif19 alters E-cadherin expression and localization. **A)** Representative confocal imaging of DMS53 cells treated with N or Kif19 siRNA that were fixed and stained for E-Cadherin and Actin. Scale bars represent 10um. **B-C)** Quantitative image analysis of individual DMS53 **(B)** and MDA-MB-231 **(C)** cells reveals that cells treated with Kif19 siRNA have significantly greater whole-cell E-cadherin levels compared to cells treated with N siRNA. Data were collected from four and five independent experiments (n=∼40 cells each) for DMS53 and MDA-MB-231, respectively. **D)** Quantitative image analysis of individual DMS53 cells expressing Myc-tagged constructs shows a significant decrease in the whole-cell E-cadherin levels of cells expressing Kif19^331-998^-Myc, but not Kif19^FL^-Myc, compared to pkMyc control. Data were collected from three independent experiments (n=∼40 cells each). **E-G)** additional analyses using the imaged cells from experiments described in graphs B-D to examine E-cadherin at cell-cell contacts. E-cadherin expression within a 1um band along cell-cell contacts was collected and corrected for band area. Data suggests that Kif19 siRNA treatment significantly increases E-cadherin at cell-cell contacts in DMS53 **(E)** and MDA-MB-231 **(F)** cells compared to N siRNA treatment. DMS53 cells expressing Kif19^FL^-Myc **(G)** have significantly decreased E-cadherin at cell-cell contacts compared to pkMyc control. Graph error bars depict mean ± standard deviation; p-values determined by Student’s t-test; ****, p<0.0001.

The role of MT dynamics at AJs is less well-defined than its role at FAs. Complete pharmacological disruption of the MT array has been shown to both enhance and disrupt e-cadherin based AJs (87, 101, 102). However, treatment with low concentrations of nocodazole, which suppresses the growth of dynamic MT plus-ends, consistently impairs AJ formation and stabilization (18, 88). Our data imply that Kif19 expression lowers both the expression of E-cadherin and its accumulation at cell-cell adhesion sites, which decreases cohesion of cells in culture. This may be related to Kif19’s role in cell invasiveness, as lower levels of E-cadherin could allow cells to more easily break away from a parent tumor and invade the surrounding environment (10, 12). Our earlier spheroid assays support this view: fewer cells were observed to migrate away from the spheroid bulk after Kif19 knockdown, which is likely secondary to increase cell-cell adhesion.

## Conclusions

We have shown that Kif19 is a potential oncogene that positively regulates cell invasiveness. Kif19 differs from other kinesin-8 family members in that it is not associated with mitosis and cell survival, which is reflected in our data showing that Kif19 knockdown does not alter cell viability. Rather, Kif19 increases the invasiveness of cancer cells, both in a 2D transwell invasion assay and a spheroid seeding assay. Our analyses indicating that Kif19 expression is higher in metastases from colorectal and breast cancer than in the primary tumor further supports a pathologic role for Kif19 in cancer. While previous studies have examined the action of an isolated Kif19 motor on MTs, our data represent the first studies to characterize MT alteration by the full-length protein, as well as the role of the CTT (64, 65). We have shown that Kif19 exhibits both MT stabilizing and destabilizing activity, and accumulates at MT plus-ends via a mechanism requiring the full-length protein. These characteristics are similar to other kinesin-8 members and are often regulated by the protein’s CTT (50, 52-54, 81). We have identified several EB-binding motifs and an NLS-sequence in the tail of Kif19, and in future studies we will use mutagenesis to delineate the full function of Kif19’s CTT (**Fig S5**) (103-106).

Kif19 was previously described as a regulator of cilia length in mice (64). Our data indicates that Kif19 expression in two unciliated human carcinoma cell lines is negatively correlated with FA density and E-cadherin expression. We propose that these changes in cell adhesion are responsible for Kif19’s effect on cell invasiveness, as decreased E-cadherin expression and changes in FA dynamics are both associated with increased cancer cell invasion (12-14, 93, 94). However, the mechanism by which Kif19 alters cell adhesion has not been defined. Our current working model is that aberrant Kif19 expression alters MT growth and stability in the cell periphery and disrupts local Rho GTPase activity. This could explain how Kif19 alters FA dynamics and the accumulation of E-cadherin at cell-cell contacts, as both can be regulated by Rho GTPases (22-25, 96). MT dynamics have been shown to locally influence the activation states of Rho GTPases, which alters actin dynamics (107). In general, MT polymerization favors Rac1 activation, as unpolymerized tubulin can sequester Rac1 and the Rac1-activator Tiam1 is activated by polymerized MTs (108, 109). This encourages actin polymerization and the initial formation of AJs and immature FAs (22, 23, 110). Conversely, MT depolymerization favors RhoA activation due to release of the RhoA-activator GEF-H1, which is sequestered and deactivated by polymerized MTs (111). RhoA promotes contraction of Myosin II, which leads to tension at FAs and AJs via actomyosin stress fibers (25, 112). This promotes the maturation of the structures, though excessive RhoA activity has also been proposed to disrupt AJs (110). The disruption of these systems, particularly the loss of myosin II-mediated tension, has been implicated in the disassembly of both FAs and AJs (95, 96, 113). In subsequent studies, we intend to use Rac1 and RhoA biosensors with FRET to examine how Rho GTPase activity is affected by Kif19 expression.

FAs and AJs are classically considered unrelated, or even competing, structures, with high FA component activation associated with E-cadherin suppression and vice-versa (7, 114, 115). Recent work has begun to reveal similarities between the two, in terms of their shared structural components, tension-mediated maturation, and regulation by the Rho GTPases (18-25). Microtubules present an additional regulatory pathway common to FAs and AJs, but there is currently a dearth of work relating all three systems. Future work on Kif19 could help elucidate a common pathway between MTs, FAs, and AJs, as well as expand our understanding of how these systems are dysregulated in cancer invasion and metastasis.

## Acknowledgements

We thank Hillary Guzik, Vera DesMarais, and, in particular, Andrea Briceno from the Einstein Analytical Imaging Facility for their training and assistance on the use of their Leica SP5 confocal microscope.

Additionally, we thank Dr. Yoshihide Hayashizaki of RIKEN OSC and Dr. Sumio Sugano of Tokyo University Graduate School for providing Kif19 cDNA from the RIKEN BioResource Center.

## Supporting Information

**S1 Fig. Kif19 expression and survival in cancer.** The R2 Genomics Analysis and Visualization Platform (http://r2.amc.nl http://r2platform.com) was used to generate Kaplan-Meier plots for individual cancer patient data sets based on Kif19 mRNA expression. The cut off for “high” vs. “low” Kif19 mRNA expression was automated by the program, and the p-values were generated using a Logrank test. Data indicate that high Kif19 expression is associated with lower overall survival in the six cancer patient datasets presented.

**S2 Fig. Kif19 knockdown and antibody validation. A)** Western blot analysis of whole cell lysates from DMS53 cells collected 72 h after treatment with control (N) or Kif19 siRNA. Blot was cut horizontally just below the 50kDa protein ladder band (denoted by black line). The top portion was probed for Kif19 (HPA073303), while the lower portion was probed for GAPDH (loading control). Densitometry measurements using Image Lab indicate a significant 61.4% decrease in Kif19 levels after siRNA treatment in DMS53 cells (n=4, Student’s t-test p<0.0001). **B)** Western blot analysis of whole cell lysates from MDA-MB-231 cells collected 72 h after treatment with N or Kif19 siRNA. Blot was cut horizontally just below the 50kDa protein ladder band (denoted by black line). The top portion was probed for Kif19 (HPA073303), while the lower portion was probed for GAPDH (loading control). Densitometry measurements using Image Lab indicate a 41.9% decrease in MDA-MB-231 cells (n=2). Yellow boxes indicate the location of the primary Kif19 transcript. Kif19 has a predicted molecular weight of 111kDa, yet we sometimes observe two bands flanking 100kDa, as is illustrated in (A). As this lower band is not always present, it is possible that it is a partial degradation product. **C)** Western blot validating binding of the Kif19 antibody primarily used in this study (Sigma Aldrich, HPA073303). Lysates are from U2OS expressing tdTomato-Kif19^FL^ or tdTomato control. The blot was cut horizontally just below the 50kDa protein ladder band and the top portion was cut vertically down the center (denoted by black lines). The left top portion was probed for Kif19, the right top portion was probed for RFP, and the bottom portion was stained for GAPDH. Note that the Kif19 band is much higher in this blot due to the added weight of the tdTomato tag. **D)** Western blot validating the binding of a second Kif19 antibody (Sigma Aldrich, AV34084). Lysates are from DMS53 cells expressing no plasmid, pkMyc control, or pkMyc-Kif19^FL^. The blot was split in half vertically (denoted by the black line). The left half was probed for Kif19 and the right was probed for Myc.

**S3 Fig. Kif19 mRNA is significantly unregulated in patient-derived metastatic cancer cells from breast and colon cancers.** Box plots showing *Kif19* mRNA expression using public DNA microarray data of tissues derived from **A)** 66 patients with primary breast carcinomas (https://www.ncbi.nlm.nih.gov/geo/query/acc.cgi?acc=gse29431), 58 breast cancer metastases samples from different organs (116), **B)** 59 primary colorectal tumor samples (117) and 24 samples of patient-derived metastatic colorectal cancer cells (117). *Kif19* mRNA is significantly unregulated in metastatic breast (p=**7.7e-03**) and colorectal cancers (p=**4.8e-09**). These plots are a useful means of presenting differences between populations as they display groups of numerical data (in this case, signal intensity/expression level of *Kif19* gene in breast and colorectal cancer tissue samples derived from human patients) through their five number summaries: the smallest observation (sample minimum= lower line), lower quartile (Q1 = bottom of box), median (Q2= line in box), upper quartile (Q3= top of the box).

**S4 Fig. Kif19-FA interactions. A)** Representative confocal imaging of MDA-MB-231 cells illustrating the extensive overlapping between adjacent FAs. Cells were transfected with tdTomato plasmid and N (left) or Kif19 (right) siRNA, then fixed 72 h later and stained for Vinculin and tdTomato. **B)** Representative confocal imaging of DMS53 cells transfected with N (top) or Kif19 (bottom) siRNA illustrating the increase in cell-cell contact observed following Kif19 siRNA treatment. **C)** Confocal imaging of DMS53 cells illustrating how the “effective” cell area was determined for use in Fig 4 G-I. The yellow outline in the rightmost image denotes the 2um border used to define each cells’ effective area. Scale bars represent 10um.

**S5 Fig. Kif19 C-Terminal Tail Domains.** Graphic depicting the theoretical EB-binding motifs and NLS domain in the Kif19 C-terminal tail. The EB-binding motifs are defined as Sx(I/L)P (104-106). The NLS domain was predicted using the online tool NLSMapper (103) and was scored as a 7.5/10 strength NLS domain.

**S1 Dataset. Minimal datasets for experiments presented in Fig 1.**

**S2 Dataset. Minimal datasets for experiments presented in Fig 4-5.**

**S3 Dataset. Minimal datasets for experiments presented in Fig 6.**

**S4 Dataset. Minimal datasets for experiments presented in Fig 7.**

